# Chronoecological interactions: Temporal niche-switching by black-striped mice after agonistic food competition with a dominant sympatric mouse species

**DOI:** 10.64898/2026.03.13.711595

**Authors:** Rafal Stryjek, Raffaele d’Isa, Michael H. Parsons, Karolina Szymańska, Katarzyna Socha, Marcin Chrzanowski, Korneliusz Kurek, Piotr Bebas

## Abstract

When novel nutrient-rich food sources become available to species sharing the same natural habitat, interspecies competition may arise, yielding insights into the ecological and social dynamics of the observed species. Here, we investigated food consumption patterns, and consequent social interactions, by two sympatric species of mice in response to a novel nutrient-rich food source. By deploying, in the mice’s natural habitat, baited video-monitored chambers, we collected, over a 5-month period, 1805 observations of food visiting by *Apodemus agrarius* and *Apodemus flavicollis*. We also documented interspecific encounters, with 86.7% of the cases showing agonism. In these interspecies agonistic encounters, *A. flavicollis* was always the initiator of agonism, attacking within 2 sec in 92.3% of the cases, and being dominant over *A. agrarius* in 84.6%. Analysis of food visiting behavior revealed that, initially, both species preferred nocturnality. However, after the interspecies fights, *A. agrarius* switched its temporal preference to diurnality, leading to temporal niche segregation between the two species and a significant reduction of interspecies encounters. Moreover, *A. agrarius* demonstrated hour-specific avoidance of *A. flavicollis*, visiting significantly less in hours with *A. flavicollis* compared to hours without. Through temporal niche switching, *A. agrarius* managed to access the food source safely, without fights. In contrast, *A. flavicollis* remained consistently nocturnal across the entire study.

Notably, our study presents the first 24h foraging actogram for free-living rodents. Moreover, while rodent interspecific competition is a well-known phenomenon, most of what we know about it comes from indirect observations. Direct observations of rodent interspecific interactions in nature are rare. Our work is the first direct (video-monitored) observation of temporal switch-inducing interspecies interactions in nature. As free-living rodents are currently considered a major model system for the study of interspecific competition, these results may offer precious insights for a better understanding of social dynamics, especially in asymmetric relationships. Furthermore, our findings highlight the significance of considering temporal dynamics in studies of interspecific interactions.

## 1. Introduction

Competition amongst mammals arises from the shared usage of limited resources, such as food, shelter and nesting sites (Berry and Jakobson, 1975). This competition plays a pivotal role in structuring ecosystems by influencing key factors including resource utilization, spatial distribution, reproductive strategies, predator-prey dynamics and evolutionary trajectories (Grant, 1972; Chesson, 1994; Chesson and Kuang, 2008; Lang and Benbow, 2013; Legault et al., 2020; Cao et al., 2025; Kanishka et al., 2025). Among mammals, rodents, exhibit a remarkable adaptability enabling them to thrive in almost any environment, from arctic or equatorial temperatures to the urbanized infrastructures of modern cities (Buckle and Smith, 2015). Rodents have a distinctive capability to adjust their behavior and physiology in response to environmental factors, such as fluctuating food availability (Hrvatin, 2020; Del Arco and Del Arco, 2025), seasonal changes in photoperiod (Butler and Zucker, 2009; Chi et al., 2025), temperature (Stryjek et al., 2021a, 2021b), moonlight conditions (d’Isa, 2025), presence of predators (Apfelbach et al., 2005) and interspecies competition (Pinter-Wollman et al., 2006; Simeonovska-Nikolova, 2007; Zhang et al., 2013; Katz et al., 2018; Viviano et al., 2022). These adjustments significantly enhance their resilience and chances of survival (Hansson, 1971; Buckle and Smith, 2015). By studying such mechanisms, researchers can better predict evolutionary mechanisms and gain insight into how exceptionally adaptive species react to environmental variability. Behavioral plasticity, widely recognized as a key survival strategy, becomes especially vital in the face of unpredictable environments (Kronfeld-Schor and Dayan, 2008; Holt, 2009). Unpredictability is particularly relevant within anthropogenically modified environments, where environmental conditions are shifting at an accelerated pace (Simon et al., 2014).

Importantly, the adaptive strategies of rodents are not just prompted by singular environmental factors (such as the availability of resources, the density of competing populations and the presence of predators; Lima and Bednekoff, 1999), but also by their interactions. For instance, in environments in which food is abundant but predators are numerous, rodents might adopt extra cautious foraging behaviors, prioritizing safety over the quantity of food collected (Hansson, 1971; Gliwicz, 1981). Conversely, in predator-loose environments with scarce resources and high density of competing populations, extra-burrow roaming and aggressive competition for food might become more frequent. As such, rodents freely living in their natural habitat represent a complex system that can serve as a particularly useful model to study mammalian behavioral adaptability to changing environments.

### 1.1 Temporal niche switching

Variations in resource availability commonly prompt competitive dynamics amongst sympatric species. This shift, when observed by researchers, has facilitated a deeper understanding of how seasonal changes and related environmental factors, such as food availability, affect interactions among species. Consequently, we now know that there are correlations between these environmental variations and the adaptive strategies that species adopt to address such changing conditions (Elton, 2001; Gliwicz and Taylor, 2002). The chronoecology (Halle and Stenseth, 2000a, 2000b) of a species refers to the timings of its interactions with its ecosystem, including both the environment and the other species. A fundamental chronoecological variable is the temporal niche. The temporal niche of a species is the period of the day in which its individuals exhibit locomotor activity (Hut et al., 2012). Temporal niche switching (TNS) is the phenomenon referring to the adjustments of activity patterns in response to changed environmental factors. For a review of TNS across phylogeny, see Hut et al., 2012.

To cope with competitive pressures, rodents have developed various foraging strategies that maximize resource access whilst minimizing interspecies conflict over food and risks of predation. These behaviors are multifaceted, encompassing dietary preferences, spatial use of the environment and TNS. In particular, when a highly sought-after food source generates strong interspecies competition, the physically inferior species may avoid this competition through niche segregation, that is by separating its niche from the that of the dominant competitor. Niche segregation can be either spatial (obtained by changing the living space) or temporal (obtained by changing the temporal use of the same living space). Interestingly, compared to the majority of the other animals, which generally rely on spatial segregation, within rodents many species adopt temporal shifting strategies. Rodent species capable of TNS include, for example, laboratory mice (Mrosovsky and Hattar, 2005), spiny mice (Pinter-Wollman et al., 2006; Cohen et al., 2010), the African striped mouse (Richardson et al., 2023), the ice rat (Oosthuizen, 2020), the tuco-tuco (Tachinardi et al., 2015) and lactating common voles (van der Vinne et al., 2014). Notably, while spatial niche changes have been extensively studied, temporal niche changes have in comparison been far less investigated, providing numerous opportunities for unexpected discoveries. For this reason, in the present work we decided to focus on temporal niches and their change (TNS) as adaptive response to competition.

The concept of temporal niche is strictly related to that of circadian rhythm, which is the daily sleep-wake cycle regulated by an organism’s internal clock. The biological clock is a complex and highly plastic system capable of synchronizing behavioral rhythms to changing environmental conditions – especially since the time-giving cues, the Zeitgebers, can be and often are biotic factors (Thoré et al., 2024). However, the study of circadian rhythms under natural conditions – including their variability during ontogeny and possible modifications – remains poorly understood, as chronobiological research has primarily focused on laboratory-based studies. In mammals, endogenous rhythms are regulated by the suprachiasmatic nucleus which drives and coordinates the timing of physiological and behavioral processes – also indirectly through peripheral clocks, which are present in nearly all organs of the body (Phillips et al., 2013; Jabbur et al., 2024).

Another related concept is diel activity, which is the distribution of an animal’s behavioral activity throughout the daily cycle (Vazquez et al., 2019; Vallejo-Vargas et al., 2022). Notably, diel activity is a broader concept than circadian rhythm, as it is determined not just by internal clocks, but also by ecological interactions with the external environment, such as when an animal eats and what it eats, which may, for instance, cause sleepiness during digestion (Hazlerigg and Tyler, 2019). TNS is a form of diel activity plasticity, which is an animal’s flexibility in its diel activity pattern (Devarajan et al., 2025).

TNS is an effective strategy in environments where resource availability fluctuates during the day or year. This strategy is sometimes used by a smaller species, such as *Mus musculus*, to avoid competition with a larger species like *Rattus norvegicus*. Muricide is a common outcome in such occurrences. Hence, TNS can be a life-saving defensive strategy for *Mus musculus*. When TNS occurs, time niche partitioning is observed, meaning that two or more species occupy different time niches of the day. Notably, the flexibility of the behavioral patterns of a species, including TNS, can offer insights into the adaptive strategies that species employs to survive in changing environments (MacArthur and Wilson, 2001; Holt, 2009; Huneman, 2019). Understanding how these strategies enable species adaptability may reveal differential capabilities among species to withstand anthropogenic pressures, climate change and sudden natural disturbances, including cataclysms (Ramírez-Bautista et al., 2020; Wan et al., 2022; Taillie et al., 2023; Hua et al., 2025).

### 1.2 Interspecies competition between black-striped mice and yellow-necked mice

The black-striped mouse (*Apodemus agrarius*) and the yellow-necked mouse (*Apodemus flavicollis*) are common in central and eastern Europe and are closely related, making them suitable subjects for comparative research (Simeonovska-Nikolova, 2007). These species offer a valuable model for investigating the mechanisms that influence patterns of activity and social behavior. Indeed, due to anthropogenic and climate-induced stochasticity, these two species occasionally co-inhabit the same peri-urban environments. Because of their overlapping habitats, agonistic interactions are frequent and competition is high among these species (Andino et al., 2011). Population density evaluations suggest that a significant overlap in food resources is the primary cause of the competition between these two species (Gliwicz, 1981). The literature reports that *A. flavicollis* is a dominant species over *A. agrarius*, with dominance frequently established through interference and aggression (Gliwicz, 1981; Mažeikytė, 2002). Additionally, laboratory studies further support this pattern: in encounters with the wood mouse (*Apodemus sylvaticus*), an *Apodemus* species similar in size to *A. agrarius*, *A. flavicollis* is clearly dominant (Hoffmeyer, 1973; Montgomery, 1978). Specifically, in encounters within a large indoor pen, *A. flavicollis* turned out to be dominant in 103 cases, while *A. sylvaticus* just in 11 cases (Hoffmeyer, 1973). Moreover, in dyadic staged encounters in a testing arena, *A. flavicollis* was dominant over *A. sylvaticus* in 63 out of 70 cases, whereas *A. sylvaticus* was dominant in only one case (Montgomery, 1978).

In our study, we evaluate the mechanisms influencing the activity patterns of *A. agrarius* and *A. flavicollis* in response to the introduction of a novel food source in a peri-urban environment. Understanding how these species respond to a novel food source and interspecific competition can enable to interpret broader ecological interactions and highlight the general adaptive strategies that allow rodents to live and thrive in changing environments (Rymer et al., 2013; Ramírez-Bautista et al., 2020; Wan et al., 2022; Taillie et al., 2023; Hua et al., 2025).

Our primary hypothesis states that either of the two sympatric species of rodents will exhibit temporal niche switching behavior when provided with a rich, novel food source. Currently, most research demonstrating niche switching involves a single species shifting from dark phase to light phase activity, as seen when the common spiny mouse changes its behavior from nocturnal to diurnal in response to light masking (Cohen et al., 2010), or when the subterranean rodent tuco-tuco switches to diurnality after spontaneous suppression of wheel-running (Tachinardi et al., 2015). On the other hand, in the present study, we observe the activity of two species concurrently, which allows to investigate the effect of interspecies social dynamics on temporal preferences. If interspecies competition is involved, we expect one of the two species to become diurnal while the other remains nocturnal. To strengthen the case for competition as the explanation for the temporal switch, our secondary experimental questions consider whether, after the introduction of the novel food source, a) most interspecific social interactions will display agonism, b) whether this will lead to the establishment of dominance hierarchies between these two sympatric rodent species, c) whether the temporal switch will be performed by the subordinate species, and d) whether the frequency of interspecific encounters will change after a temporal switch (i.e., whether the temporal switch is actually functional in obtaining avoidance of the dominant competitor).

Importantly, ecological niche segregation in response to interspecies competition is a well-known phenomenon among rodents (Grant, 1972; Eccard and Ylönen, 2003). Nevertheless, up to date, this phenomenon has never been observed directly. Notably, the present study is the first direct (video-monitored) observation of temporal switch-inducing interspecies interactions in nature. Moreover, while complete daily actograms for rodents are commonly recorded in the laboratory, in the wild they are rarely recorded and they are mostly based on generic motion data from biologgers (for example, Hoogenboom et al., 1984; Šklíba et al., 2007, 2014, 2025; Lövy et al., 2013; Silvério and Tachinardi, 2020; Silvério et al., 2025). Here, we present the first 24h foraging actogram for free-living rodents. Remarkably, the so called "wild clocks" approach (Oda and Valentinuzzi, 2024) is particularly relevant, as it has been observed several times that rodents changed their biological rhythms after having been transported to the laboratory, for example switching from diurnal to nocturnal activity (Blanchong and Smale, 2000; Hagenauer and Lee, 2008; Kronfeld-Schor et al., 2013; Fritzsche et al., 2017). Hence, the study of rodent biological rhythms just in the laboratory may lead to findings that do not actually have ethological validity. Our approach can contribute to deepen our ethologically and ecologically relevant knowledge of rodent temporal preferences.

## 2. Material and Methods

### 2.1 Studied species

In this study, two rodent species were observed, the black-striped mouse (*Apodemus agrarius*) and the yellow-necked mouse (*Apodemus flavicollis*). Photos are shown in Figure 1A and 1B. These species share a common habitat from Central Europe to the Urals. The former is common mainly in grasslands, agricultural lands, and also at the edges of forests. It is also known for being interspersed and adapting to various environmental conditions, including urban environments (Gliwicz, 1981; Łopucki and Kiersztyn, 2020). The latter has been defined a forest specialist (Gasperini et al., 2025), as it prefers mixed forests, where it plays a vital role in seed dispersion as well as in aeration of the soil (Juškaitis, 2002). However, in areas at the borders of forests, *A. agrarius* and *A. flavicollis* share the same environments. Because of the overlapping habitats of these two species, we expect frequent agonistic interactions and competition. As a separate part of this study, recently published (d’Isa et al., 2024a), we observed that *A. agrarius* engages in an intricate and deceptive game of hide and seek with *A. flavicollis*. This peculiar avoidance behavior from *A. agrarius* in response to interspecies antagonism could relate to the fact that it is endowed of a higher level of behavioral plasticity, while *A. flavicollis* has superior physical adaptive modifications (Gliwicz, 1981).

*A. agrarius* can be distinguished by several morphological features.The upper parts of *A. agrarius* are covered with grayish-brown fur with a rusty tone, and a black stripe runs through the body, giving the species its common name. The underparts of the mouse are paler and gray. Ears and eyes of *A. agrarius*, in comparison with representatives of other species of rodents, are rather small, which could be associated with its particular sensory perception and behavioral features (Balčiauskas et al., 2021).

*A. flavicollis* is on average bigger than *A. agrarius*. The top parts of the *A. flavicollis* are a distinctive brownish-grey color, which provides camouflage in its natural habitat. The underparts are white. A characteristic feature of this species is the yellow band of fur around the neck. Juvenile yellow-necked mice have the dorsal fur of a paler tone of greyish brown when compared with adults, indicating that individuals of this species undergo a modification in fur coloration as they mature.

**Figure 1.**
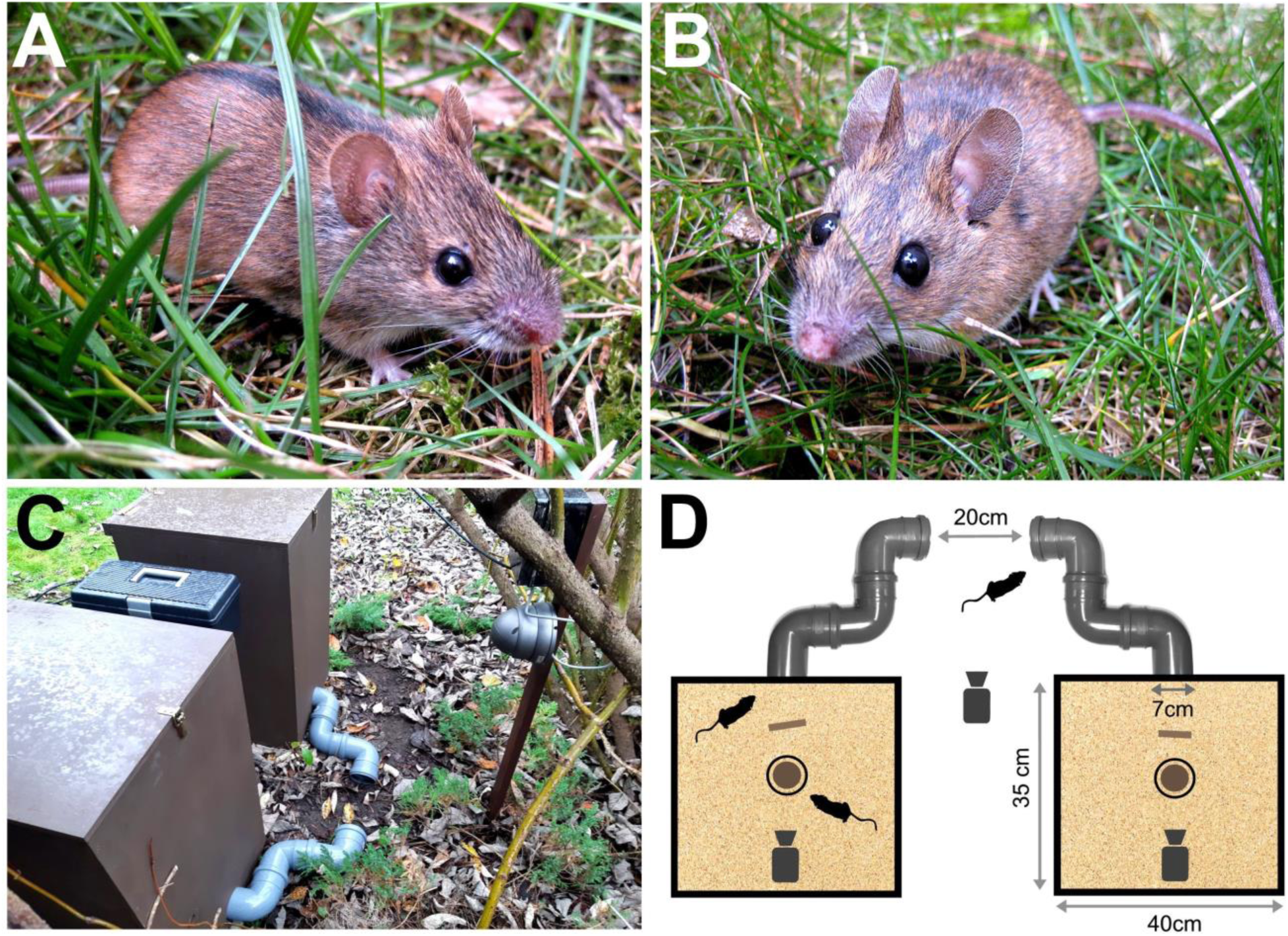
Monitored species and test equipment. A: Black-striped mouse (*Apodemus agrarius*). B: Yellow-necked mouse (*Apodemus flavicollis*). C: External view of the test chambers. D: Schematic representation of the test chambers. Motion-activated video cameras record the behavior of free-ranging mice outside and inside the test chambers. The brown circle at the chamber’s center represents the bait (chocolate-nut cream). The brown rectangle represents a piece of wood. The floors of the chambers are covered with sand.

Notably, our team has sampled *Apodemus* mice from the same location of the present study, finding that male *A. flavicollis* mice were, on average, considerably bigger than male *A. agrarius* mice. These mice have been collected by our team for independent biochemical analyses that in part have been published (Stryjek et al., 2024) and in part are upcoming (Stryjek et al., in preparation). We are reporting here (Supplementary Figure 1) the morphometric data for male *A. agrarius* and *A. flavicollis* mice sampled from the location of the present study, more specifically body length, total length (body + tail length) and weight. Remarkably, *A. flavicollis* was largely superior to *A. agrarius* in all three morphometric variables. In particular, the average total body length was 143 mm for *A. agrarius* and 192.8 mm for *A. flavicollis* (+34.8% in *A. flavicollis*; p = 0.0003), with the average body length being 79.9 mm for *A. agrarius* and 95.8 mm for *A. flavicollis* (+19.9% in *A. flavicollis*; p = 0.015). The average weight was 19.5 g for *A. agrarius* and 31.7 g for *A. flavicollis* (+63% in *A. flavicollis*; p = 0.009). While *A. agrarius* reached a maximal total length of 156 mm, *A. flavicollis* reached 224 mm (+43.6%). Analogously, maximal weight was 24.6 g for *A. agrarius* and 41.8 g for *A. flavicollis* (+69.9%).

### 2.2 Study site

This study took place from November 2020 to April 2021 in our dedicated laboratory-to-field study site (Stryjek et al., 2021a, 2021b, 2024; Parsons et al., 2023a; d’Isa et al., 2024a): a peri-urban area of Warsaw (Poland) next to a forest (52°20′20.00′′N; 21°03′30.00′′E; altitude of 80 m). Test chambers were set up on private land and their deployment was authorized by the owners of the properties. Temperatures ranged from +16 °C to −20 °C. This observational study was essentially non-invasive and based on the surveillance of free-ranging animals. Animals were not marked nor handled in any way in order to avoid handling-induced stress and captive behavior. Rodents were free to enter or ignore the test chambers that were set up with food sources and video cameras. The employment of a 24-hour surveillance by motion-triggered video recording provided a full-time monitoring that allowed us to capture activity patterns, foraging behavior, and social interactions of the two species.

### 2.3 Field observation chambers

We have long utilized laboratory-style chambers, with approaches validated inside the laboratory, to extend behavioral observations to wild populations of rodents, incorporating in this way naturalistic conditions involving multiple species and contexts (Stryjek et al., 2021a, 2021b, 2024; Parsons et al., 2023a, 2023b; d’Isa et al., 2024a, 2024b). The observation chambers (Figs. 1C and 1D) were wooden boxes with 35 × 40 cm floors and 70 cm high walls. These boxes were built with 12 mm thick waterproof plywood panels painted with odorless acrylic paint purchased from Luxens, Leroy Merlin, France.

Chambers were connected to 7 cm diameter and 50 cm long plastic sewer pipes which served as a single point of entrance or exit. Two test chambers, located 20 cm apart, were used in this study. Test chambers were free-access and could be visited by free-ranging mice at any time of the day. An infrared video camera was placed inside, and outside each chamber. To enable remote monitoring, the 3 infrared Easycam EC-116-SCH; Naples, FL, United States) were connected to a digital video recorder (Easycam EC-7804 T; Naples, FL, United States). This equipment allowed animal detection and subsequent recording at all hours for the whole duration of the study. Source of food was supplied in the evenings, shortly after dusk since this time period is known to reflect the peak of nocturnal rodent activity (Stryjek et al., 2013) Ten grams of chocolate-nut cream (Nutella, Alba, Italy) was used as bait on a daily basis. The chocolate-nut cream was applied evenly on the surface of a 70 mm glass Petri dish which was placed in the middle of the chamber. The floor of each chamber was covered with 1 cm of rinsed sand and replaced after every 2 −4 days. To eliminate possible scent markings, the entrance pipes were thoroughly cleaned with the unscented liquid soap Biały Jeleń (Pollena, Ostrzeszów, Poland) every 2-4 days.

### 2.4 Video recording and behavioral scoring

The camera-equipped chambers were deployed on the evening of 1st November 2020 and the experimental observations ranged from 2nd November 2020 to 9th April 2021. Excluding 23 days with missing recordings due to technical (e.g. power failure) or meteorological (e.g. heavy snowfall) issues, we obtained a total of 136 recorded days. In order to allow an observation of trends and shifts without the data becoming too fragmented and statistically insignificant, we divided these 136 recorded days into 4 periods. In particular, to identify changes in behavior over the entire period of the study, we segregated 4 equally sized time-bins of 34 days. Exact period timeframes were as follows:

Period 1: 2020-11-03 to 2020-12-07. Habituation/Baseline: This initial period provides baseline data on the activity patterns of the species as they acclimatized to the test chambers and to the introduction of a novel food source. Period 2: 2020-12-08 to 2021-01-17. This period captures behavior in the winter season, when colder temperatures and reduced daylight could potentially influence the behavior and activity patterns of the two species. Period 3: 2021-01-18 to 2021-03-06. Period 4: 2021-03-07 to 2021-04-09. The final period of the study covered the late winter to early spring seasons. This is a time when both species might exhibit increased activity due to the approaching breeding season.

Cameras in both chambers, and immediately outside the chambers, recorded behaviors and species interactions. Only cameras inside the chambers were used for systematic analysis. The collected data included the date and time when a mouse entered or left the chamber, the chamber ID, the species of the mouse that entered the chamber, the type of interaction between the mice in the chamber (non-agonistic; agonistic: chases or fights), the initiator and target of any agonistic interaction, and the winner of any agonistic interaction (defined as the mouse that managed to make the other one escape from the chamber).

The study was conducted in different months (and thus under varying daytime lengths). To have a uniform definition of night and day, we elected to use, as the times when daytime began and ended, the civil twilights, that is the time just before sunrise (civil dawn) and just after sunset (civil dusk), when the sun is less than six degrees below the horizon. Civil twilight times were obtained from the website timeanddate (https://www.timeanddate.com/sun/poland/warsaw) by Time and Date AS (Stavanger, Norway). In this way, we were able to determine, across days with different lengths, whether a given behavior occurred during the daytime or at night., For each visit, the dark/light phase of the time of entry in the chamber (as defined by the civil twilights) was reported. This approach allowed us to standardize the collected information and detect possible switches in the temporal niches of the studied species.

The recordings were scored by two researchers separately, by watching the videos and filling out the scoring sheet. Examples of behavioral events are reported in Figure 2. Individual visiting by *A. agrarius* and *A. flavicollis* can be seen in Figures 2A and 2B, respectively. Cases of non-agonistic and agonistic interspecies encounter are shown in Figures 2C and 2D, respectively.

**Figure 2.**
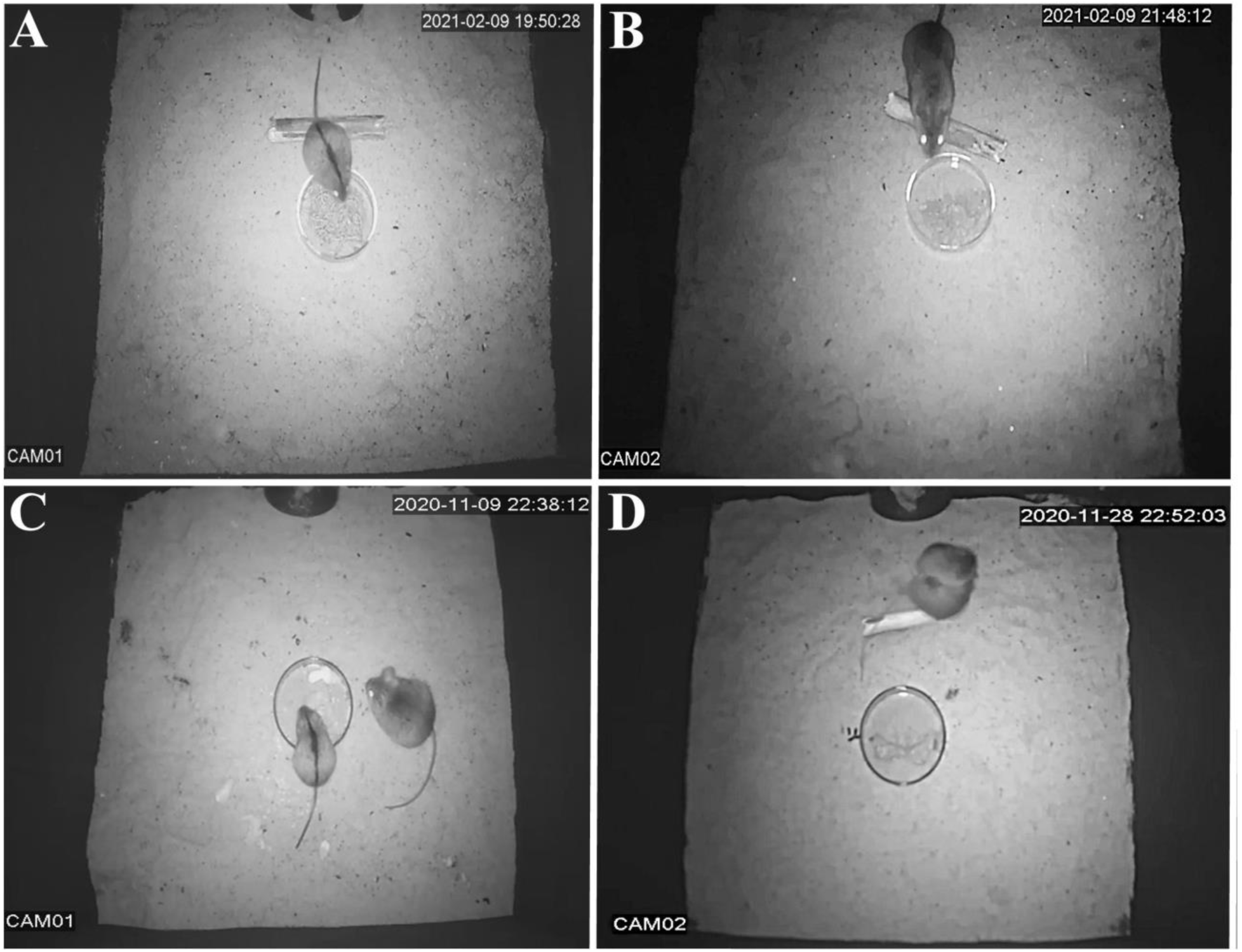
Individual chamber visiting and in-chamber social encounters. A: Individual visit by a black-striped mouse that entered test chamber 1 at 22:42:07 on the 20th of November 2020. B: Individual visit by a yellow-necked mouse that entered test chamber 2 at 20:50:54 on the 8th of December 2020. C: Non-agonistic social interaction between the two species in test chamber 1 at 22:38:12 on the 9th of November 2020. D: Agonistic social interaction between the two species in test chamber 2 at 22:52:03 on the 28th of November 2020.

### 2.5 Statistical analysis

For the analysis of temporal preferences in relation to time of the day, we first counted the number of visits occurring in each ofthe 24 hours of the day. From these values, we calculated the percentage of activity for each hour: percentage of activity for hour X = (number of visits in hour X / number of visits in all 24 hours) x 100. Hourly percentages of activity were compared between species through two-proportion z-tests. This analysis was performed for each of the 4 time periods.

For temporal preferences in relation to the light/dark phase, we first counted the number of visits occurring in the light phase (daytime) and in the dark phase (nighttime). Daytime was defined as the phase between civil dawn and civil dusk, while nighttime was defined as the phase between civil dusk and civil dawn. We then calculated the percentage of activity for each phase: percentage of activity for phase X = (number of visits in phase X / number of visits in both phases) x 100. As the percentage of daytime activity and the percentage of nighttime activity are perfectly complementary, for the subsequent analysis we chose only one of the two variables: the percentage of nighttime activity. In order to understand the chronotype of each species, for each species we compared the percentage of nighttime activity against chance level (which was obtained from the percentage of time occupied by nighttime in the analyzed period). Percentages of nighttime activity significantly higher than chance level indicate a significant preference for the dark phase (nocturnality), while percentages significantly lower than chance reveal a significant preference for the light phase (diurnality). Percentages that do not differ from chance level show indifference between the dark phase and the light (cathemerality). The within species a chance level analysis was performed through a one-proportion z-tests against chance level. Moreover, the percentages of nighttime activity were compared between species though two-proportion z-tests. The dark-light phase within-species and between-species analyses were performed for each of the 4 time periods.

For the general analysis of agonism, we classified each social encounter as non-agonistic or agonistic. Agonism was defined as a physical attack and/or a chase by one mouse towards another mouse. Social encounters were further classified, based on the identity of its members, into: interspecific, all intraspecific, *A. agrarius* intraspecific and *A. flavicollis* intraspecific. For each category, the percentage of agonism was calculated: percentage of agonism for category X = (number of agonistic encounters in category X / number of all social encounters in category X) x 100. The percentage of interspecific agonism was compared the percentages of agonism of the other categories through two-proportion z-tests.

For the analysis of interspecies aggressiveness, we scored, for each interspecific agonistic encounter, the initiator of agonism. For each species, the percentage of interspecies agonism initiation was calculated: percentage of interspecies agonism initiation by species X = (number of cases of agonistic initiation by species X / number of all interspecies agonistic encounters) x 100. Percentages of interspecies initiation were compared against chance level. As all interspecies encounters were dyadic, such chance level was 50%.

For the analysis of interspecies dominance, we scored, for each interspecific agonistic encounter, the winner of the agonistic interaction (the mouse that managed to make the other one leave the chamber). For each species, the percentage of dominance in interspecies agonism encounters was calculated: percentage of interspecies agonism dominance of species X = (number of cases of win by species X in an interspecies agonistic encounter / number of all interspecies agonistic encounters) x 100. Percentages of interspecies dominance were compared against chance level. As all interspecies encounters were dyadic, such chance level was 50%.

For the analysis of the changes the occurrence of interspecies encounters over the 4 time periods, we calculated how interspecies encounters were distributed across the time periods: percentage of interspecies encounters in period X = number of interspecies encounters in period X / number of interspecies encounters in all periods) x 100. Percentages were compared against chance through a one-proportion z-tests against chance level. Since the 4 periods were equally sized, such chance level was 25%. Furthermore, a direct analysis of the change of percentages of interspecies encounters across time periods was performed. McNemar’s tests for related proportion were used to compare the baseline percentage of P1 against the percentages of the following periods.

For the analysis of interspecies avoidance, the percentage of nocturnal activity of each species was compared across time periods. McNemar’s tests for related proportion were used to compare the baseline percentage of P1 against the percentages of the following periods. Moreover, for each time period, we counted the number of chamber visits by *A. agrarius* in the hours when *A. flavicollis* was present and in the hours when *A. flavicollis* was absent. For each period, the means of the *A. agrarius* visits in the *A. flavicollis*+ and in *A. flavicollis-* were compared through independent-samples t-tests.

Finally, for the analysis of the environmental effects of the arrival of the breeding season, we calculated the mean daily daytime length and mean daily maximal temperature for the period before the breeding season (P3) and for the period of the breeding season (P4). These means were compared through independent-samples t-tests. The website timeanddate served as source for the daytime lengths (https://www.timeanddate.com/sun/poland/warsaw) and the temperatures (https://www.timeanddate.com/weather/poland/warsaw).

Statistical analyses were performed using the following software: IBM SPSS Statistics 23.0, MedCalc 23.4 and Social Science Statistics. Statistical significance was set at p < 0.05. For the visualization of the significant differences, a 4 stars system was employed: * p < 0.05, ** p < 0.01, *** p < 0.001 and **** p < 0.0001.

## 3. Results

Chamber visiting behavior of free-living *Apodemus* mice was recorded over the course of 136 days, subdivided into 4 equally sized periods of 34 days each (P1-P4). In the entire study, we recorded a total number of 1805 visits in the chambers, by *A. agrarius* and *A. flavicollis*. Both species discovered and entered the chambers within 40h from their deployment, and continued to show chamber visiting throughout all four time periods, up to the last day of the study. Nevertheless, the temporal pattern of activity notably differed between the two sympatric species, indicating diversified temporal niches.

### 3.1 Temporal preferences in relation to time of the day

In order to evaluate the temporal preferences of the two species, firstly we analyzed the visits in relation to the time of the day. Figure 3 shows the distribution of the activity of the two species across the 24 hours of the day, from the first (00:00-00:59) to the last (23:00-23:59). Across the daily hours, the two species exhibited very different temporal preferences. In particular, in the first period (P1) the activity of *A. agrarius* was significantly higher than the one of *A. flavicollis* at all hours between 7 and 19 (8^th^ h: z = 2.383, p = 0.017; 9^th^ h: z = 2.355, p = 0.018; 10^th^ h: z = 2.034, p = 0.042; 11^th^ h: z = 2.355, p = 0.018; 12^th^ h: z = 2.729, p = 0.006; 13^th^ h: z = 3.292, p = 0.00099; 14^th^ h: z = 3.292, p = 0.00099; 15^th^ h: z = 2.355, p = 0.018; 16^th^ h: z = 2.146, p = 0.032; 17^th^ h: z = 3.784, p = 0.0002; 18^th^ h: z = 3.580, p = 0.0003; 19^th^ h: z = 3.284, p = 0.0010). On the other hand, the activity of *A. flavicollis* was significantly higher than the one of *A. agrarius* at all hours between 23 and 6 (24^th^ h: z = 2.526, p = 0.011; 1^st^ h: z = 3.544, p = 0.0004; 2^nd^ h: z = 5.698, p < 0.0001; 3^rd^ h: z = 4.625, p < 0.0001; 4^th^ h: z = 2.786, p = 0.005; 5^th^ h: z = 4.506, p < 0.0001; 6^th^ h: z = 3.227, p = 0.0012). At the hours between 19 and 23, the activities of the two species were statistically comparable.

**Figure 3.**
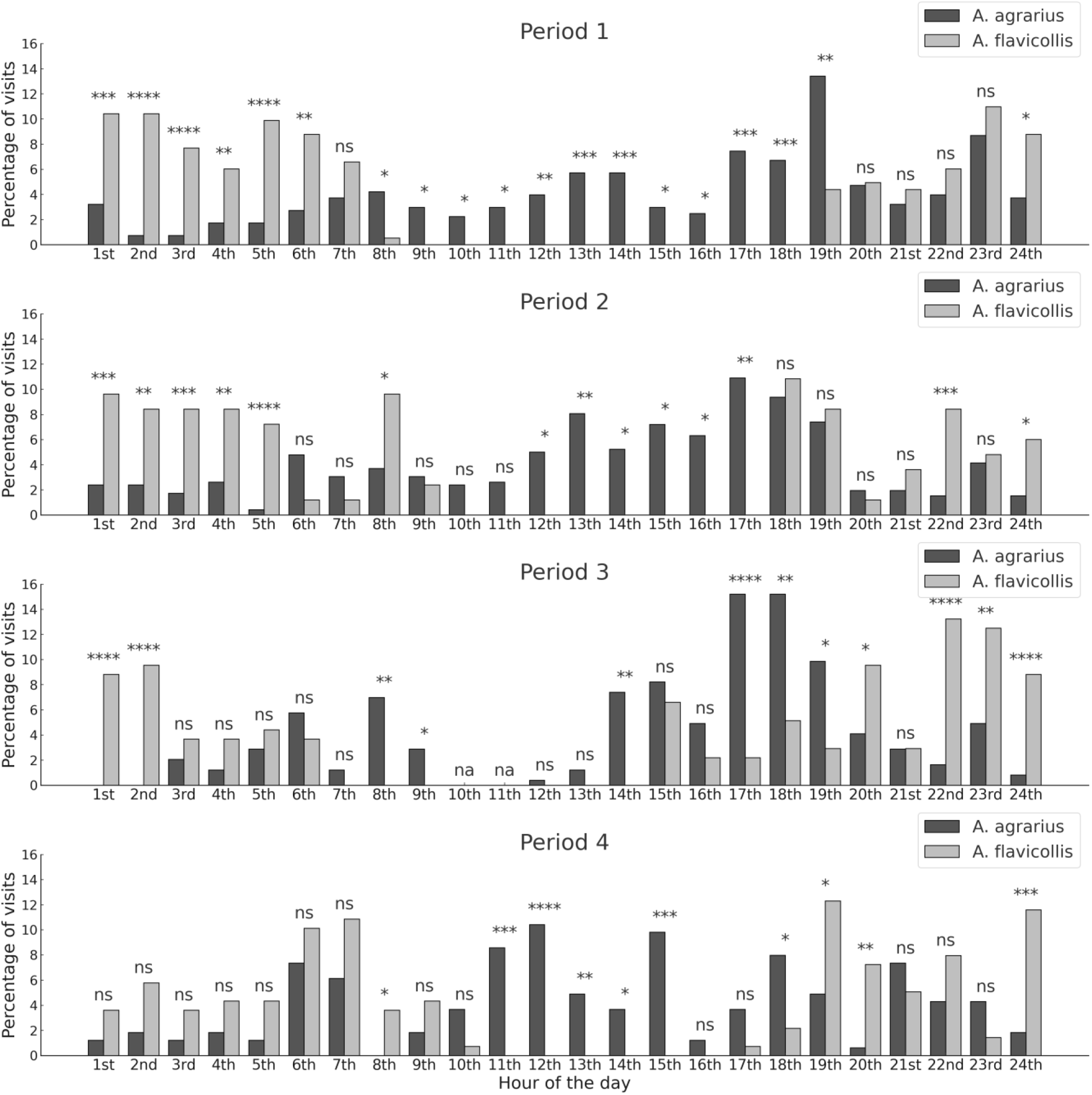
Temporal preferences of *A. agrariu*s and *A. flavicollis* in relation to time of the day. The actograms show the percentage of visits for the 24 hours of the day, from the 1st (00:00-00:59) to the last (23:00-23:59), across the 4 periods of the study (P1-P4, from top to bottom). Asterisks highlight significant differences between species (* p < 0.05, ** p < 0.01, *** p < 0.001 and **** p < 0.0001; ns: not significant).

In P2, the activity of *A. agrarius* was significantly higher than the one of *A. flavicollis* from 11 to 17 (12^th^ h: z = 2.086, p = 0.037; 13^th^ h: z = 2.683, p = 0.007; 14^th^ h: z = 2.133, p = 0.033; 15^th^ h: z = 2.524, p = 0.012; 16^th^ h: z = 2.357, p = 0.018; 17^th^ h: z = 3.160, p = 0.002), while the activity of *A. flavicollis* was greater from 23 to 5 (24^th^ h: z = 2.559, p = 0.010; 1^st^ h: z = 3.295, p = 0.00096; 2^nd^ h: z = 2.819, p = 0.005; 3^rd^ h: z = 3.414, p = 0.0006; 4^th^ h: z = 2.647, p = 0.008; 5^th^ h: z = 4.717, p < 0.0001), as well as at the 8^th^ h (z = 2.366, p = 0.018) and 22^nd^ h (z = 3.646, p = 0.0003).

In P3, the activity of *A. agrarius* was significantly higher from 7 to 9 (8^th^ h: z =3.156, p = 0.002; 9^th^ h: z = 1.998, p = 0.046), at the 14^th^ h (z = 3.252, p = 0.0012) and from 16 to 19 (17^th^ h: z = 3.957, p < 0.0001; 18^th^ h: z = 2.938, p = 0.003; 19^th^ h: z = 2.476, p = 0.013). Conversely, the activity of *A. flavicollis* was significantly greater than the one of *A. agrarius* from 21 to 2 (22^nd^ h: z = 4.628, p < 0.0001; 23^rd^ h: z = 2.656, p = 0.008; 24^th^ h: z = 3.961, p < 0.0001; 1^st^ h: z = 4.706, p < 0.0001; 2^nd^ h: z = 4.904, p < 0.0001) and at the 20^th^ h (z = 2.129, p = 0.033).

Finally, in P4, the activity of *A. agrarius* was significantly higher than the one of *A. flavicollis* from 10 to 15 (11^th^ h: z = 3.526, p = 0.0004; 12^th^ h: z = 3.906, p < 0.0001; 13^th^ h: z = 2.638, p = 0.008; 14^th^ h: z = 2.277, p = 0.023; 15^th^ h: z = 3.782, p = 0.0002) and at the 18^th^ h (z = 2.236, p = 0.025). On the other hand, *A. flavicollis* was significantly more active than *A. agrarius* from 18 to 20 (19^th^ h: z = 2.322, p = 0.020; 20^th^ h: z = 3.056, p = 0.002), at the 24^th^ h (z = 3.467, p = 0.0005) and at the 8^th^ h (z = 2.451, p = 0.014).

### 3.2 Temporal preferences in relation to the light/dark phase

In order to obtain a more ecologically relevant evaluation of the diel phenotypes of the two species, we then analyzed their activity in relation to the light/dark phase. Based on light conditions, animal temporal activity can be classified into diurnal (preferentially distributed in the daytime), nocturnal (preferentially distributed in the nighttime) or cathemeral (indifferently distributed across daytime and nighttime) (Halle, 2006; Bennie et al., 2014; Vallejo-Vargas et al., 2022; Devarajan et al., 2025). To evaluate the preferences for the light or the dark phases, we compared, for each time period, the percentage of activity in the dark phase against chance level (which is obtained from the percentage of time occupied by the dark phase in the given period). Percentages significantly higher than chance level indicate a significant preference for the dark phase (nocturnality), while percentages significantly lower than chance demonstrate a significant preference for the light phase (diurnality). Percentages that do not differ from chance level show indifference between the dark and the light phases (cathemerality). In our study, the two species exhibited different light-related temporal niche preferences.

Percentages of nighttime activity across the four periods are shown in Figure 4A. Interspecies analysis revealed that *A. flavicollis* exhibited significantly more nocturnal activity than *A. agrarius* in all 4 time periods (P1: z = 7.656, p < 0.0001; P2: z = 5.769, p < 0.0001; P3: z = 7.547, p < 0.0001; P4: 6.819, p < 0.0001). Notably, in every time period, *A. flavicollis* had percentages of nocturnal activity that were more than 30% over the ones of *A. agrarius* (P1: +30.1%; P2: +33.8%; P3: +38.8%; P4: +39.4%).

**Figure 4.**
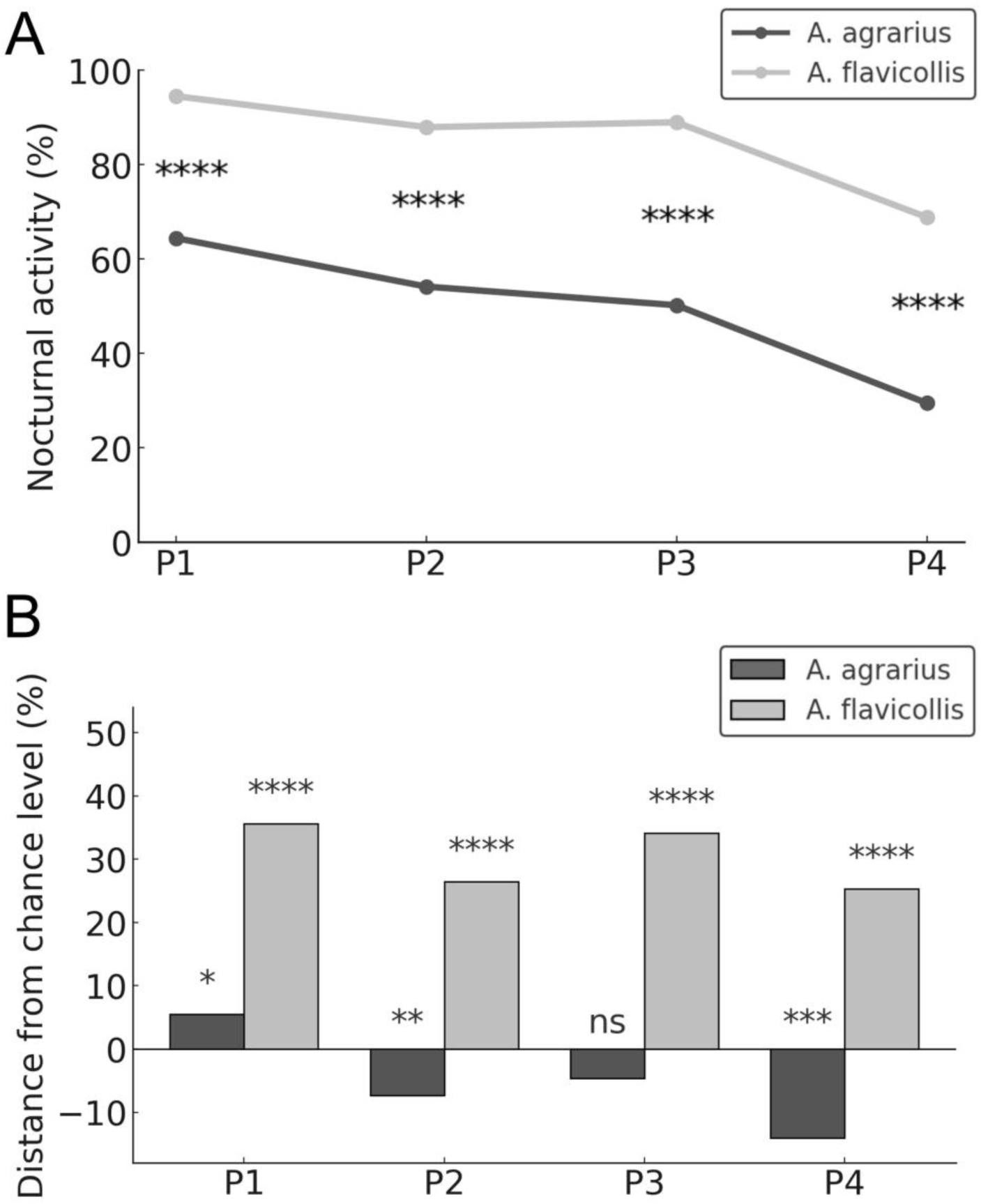
Temporal preferences of *A. agrariu*s and *A. flavicollis* in relation to the dark/light phase. A: Percentage of nighttime visiting across the 4 time periods P1-P4. Asterisks highlight significant differences between species (**** p < 0.0001). B: Distance between the observed percentage of nighttime visiting and the chance level percentage of nocturnal visiting for each period. Asterisks highlight significant differences against chancel level (* p < 0.05, ** p < 0.01, *** p < 0.001 and **** p < 0.0001; ns: not significant).

The chance level analysis clarified that, over the course of the study, the two species had different chronotypes. Figure 4B illustrates the distance from chance level for the nocturnal activity of each species across the 4 time periods. Significantly positive distances indicate nocturnality, while significantly negative distances indicate diurnality. Percentages at chance level demonstrate cathemerality.

In the initial period (P1), *A. agrarius* exhibited nocturnality. The percentage of nighttime visiting was significantly higher than chance level (distance from chance level: +5.5%; z = 2.254, p = 0.024). However, in P2, *A. agrarius* switched to diurnality, with the percentage of nighttime visiting being significantly lower than chance level (distance from chance level: −7.3%; z = −3.230, p = 0.0012). In P3, the percentage of nighttime visiting of *A. agrarius* was not significantly different from chance level (z = 1.457, p = 0.145), indicating cathemerality. Finally, in P4 a strong shift towards diurnality was observed again, with a percentage of nighttime visiting significantly below chance level (distance from chance level: −14.1%; z = −3.622, p = 0.0003).

On the other hand, *A. flavicollis* showed a very strong nocturnality in all time periods. In P1, almost the entire activity of *A. flavicollis* was nocturnal (94.5%). The percentage of nighttime visiting was significantly higher than chance level (distance from chance level: +35.6%; z = 9.763, p < 0.0001). This temporal preference for nighttime was maintained in all subsequent periods: P2 (distance from chance level: +26.5%; z = 4.954, p < 0.0001), P3 (distance from chance level: +34.1%; z = 7.994, p < 0.0001) and P4 (distance from chance level: +25.3%; z = 6.002, p < 0.0001).

### 3.3 Social encounters, agonism and temporal preferences

In the entire study, we recorded 100 social encounters. Of these, almost all (98) were dyadic encounters, while 2 were triadic. The social encounters involved a total of 202 *A. agrarius* and *A. flavicollis* mice. Both intraspecies and interspecies encounters occurred, with intraspecies encounters representing the majority of the encounters (85%). In order to better understand the social dynamics, we scored the presence of agonistic behavior in both intraspecies and interspecies encounters (Figure 5A). Notably, agonism was significantly higher in interspecies encounters than in intraspecies encounters (86.7% vs 57.6%; z = 2.135, p = 0.033). The frequency of interspecies agonism was significantly higher than both the frequency of *A. agrarius* intraspecies encounters (59.2%; z = 2.021, p = 0.043) and the frequency of *A. flavicollis* intraspecies encounters (44.4%; z = 2.203, p = 0.028). The two triadic encounters featured exclusively *A. agrarius* mice, and were both non-agonistic.

**Figure 5.**
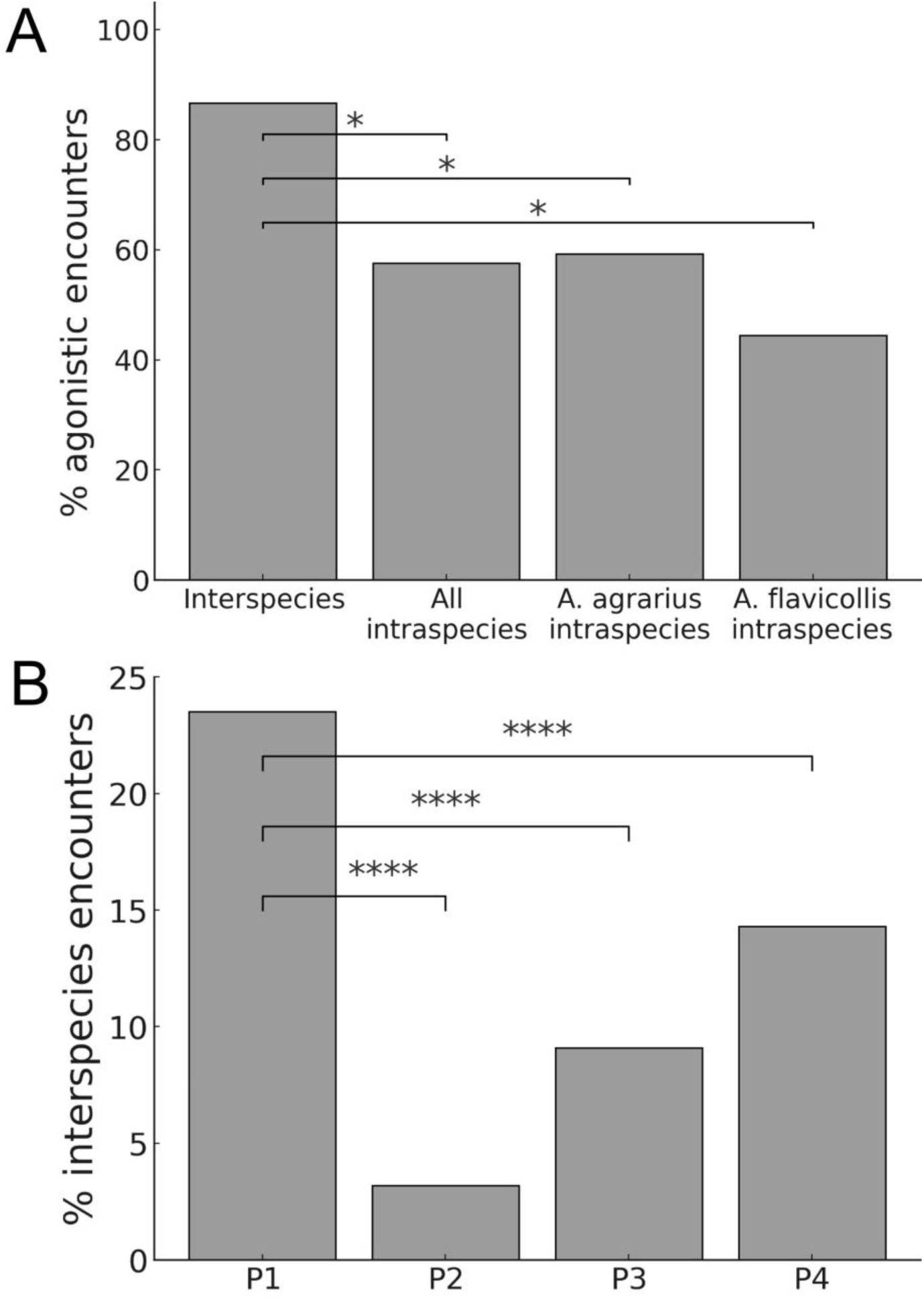
Interspecies agonism and period-related evolution of the frequency of interspecies encounters. A: Percentage of agonistic encounters in interspecies and intraspecies encounters. Asterisks highlight significant differences against interspecies agonism (* p < 0.05). B: Percentage of interspecies encounters across the 4 time periods. Asterisks highlight significant differences against the initial period P1 (**** p < 0.0001).

Remarkably, in the agonistic interspecies encounters, *A. flavicollis* was more aggressive than *A. agrarius*, and dominant over it. Aggressiveness was evaluated by scoring the initiator of the agonistic interaction. Strikingly, we observed that *A. flavicollis* was the initiator of the agonism in all interspecies encounters (z-test vs chance level: z = 3.606, p = 0.0003). Its latency to attack was impressively low, as in the vast majority of the cases (92.3%) it attacked *A. agrarius* within 2 sec from the beginning of the encounter. Moreover, *A. flavicollis* was the winner of the agonistic interaction (the mouse that managed to make the other one leave the chamber) in 84.6% of the cases (z-test vs chance level: z = 2.496, p = 0.013), indicating a very strong dominance over *A. agrarius*.

Most interspecies encounters occurred in P1 (80%), dramatically dropping down to 6.7% in P2 and remaining at that level in P3 (6.7%) and P4 (6.7%). This distribution of the interspecies encounters across the 4 equally sized time periods was highly skewed towards P1 and the percentage of interspecies encounters occurring in P1 was significantly higher than the expected chance level of 25% (z = 4.919, p < 0.0001). In order to directly compare the percentages of interspecies encounters across time periods (Figure 5B), we used McNemar’s test for related proportions. In P1, interspecies encounters were 23.5% of all encounters. In P2, the percentage of interspecies encounters dramatically dropped to 3.2% (McNemar’s test: p < 0.0001), remaining at significantly lower levels in all remaining time periods, P3 (9.1%; McNemar’s test: p < 0.0001) and P4 (14.3%; McNemar’s test: p < 0.0001).

Importantly, this drop in the frequencies of interspecies encounters after P1, appears to be due not to a change in the behavior of *A. flavicollis*, but to a change in the behavior of *A. agrarius*. Indeed, *A. flavicollis* maintained a strong preference for nocturnality across all 4 time periods, with its percentages of nocturnal activity being more than 25% over the chance level in all time periods. On the other hand, *A. agrarius* began as nocturnal in P1 but, after the frequent interspecies fights of P1 (the interspecies encounters of P1 were agonistic in 83.3% of the cases), in P2 it then switched to diurnality, leading to a segregation of the temporal niches of the two species. In particular, *A. agrarius* changed its temporal niche, inverting its dark/light period preference and shifting its percentage of nocturnal activity from +5.5% above chance level to −7.3% under chance level. Direct comparison of visiting activity over time periods, showed that *A. agrarius* significantly reduced its percentage of nocturnal visits in P2 vs P1 (McNemar’s test: p < 0.0001). Interestingly, analysis of the visits per hour of the day further supports the idea that *A. agrarius* was avoiding *A. flavicollis* (Figure 6). Indeed, in P1 *A. agrarius* showed comparable levels of activity in the hours when *A. flavicollis* was active vs the hours when *A. flavicollis* was inactive, as revealed by the comparison of the mean number of visits in *A.* flavicollis+ vs *A. flavicollis*-hours (t = −0.235, p = 0.816). On the other hand, in P2 *A. agrarius* performed significantly less visits in the hours when *A. flavicollis* was active than in the hours when *A. flavicollis* was inactive (t = −2.524, p = 0.019).

**Figure 6.**
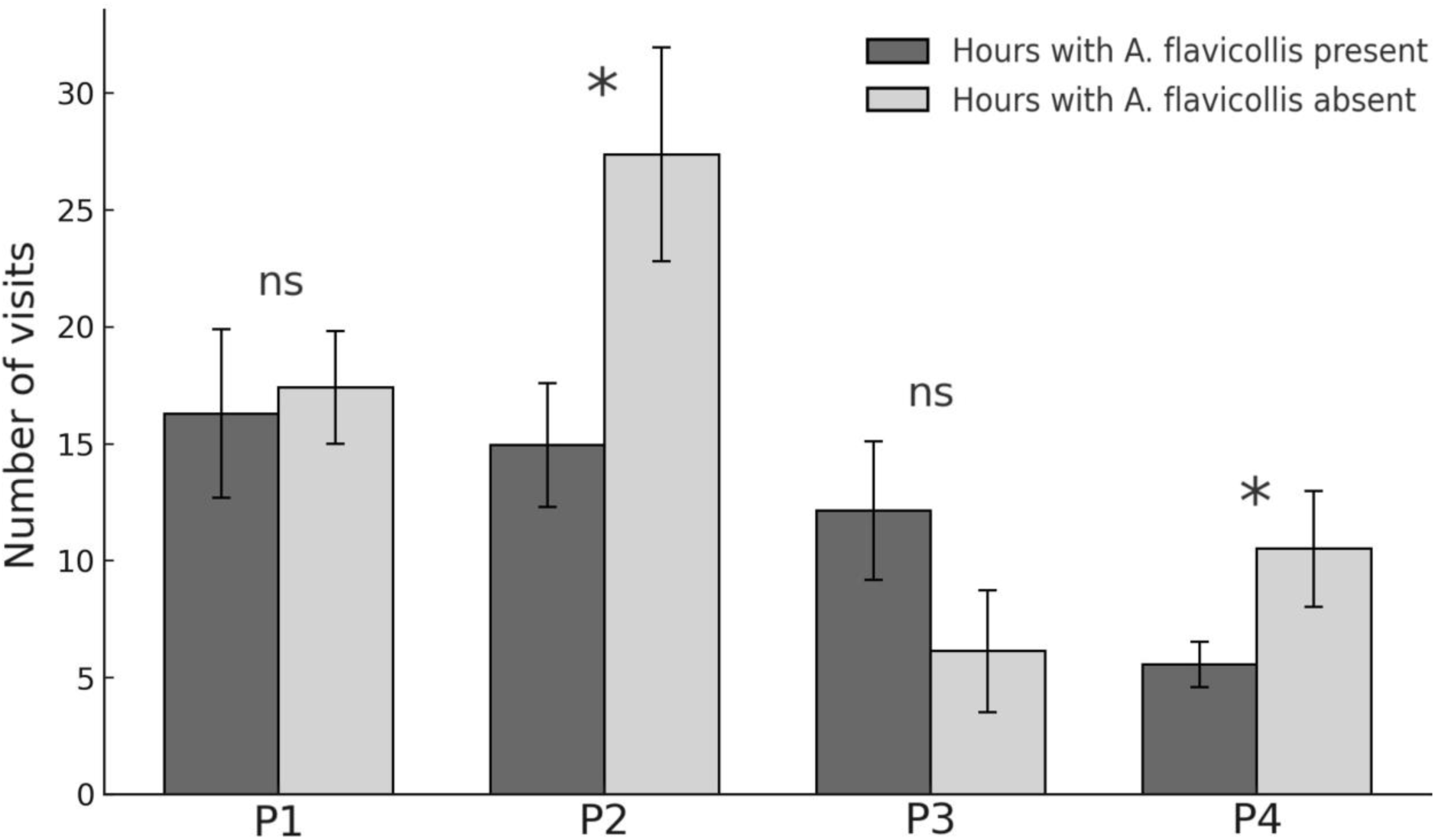
Interspecies avoidance by *A. agrarius* towards *A. flavicollis*. The graph shows the number of food visits by *A. agrarius* in the hours with and without *A. flavicollis*, for each time period. Asterisks highlight significant differences between conditions (* p < 0.05; ns: not significant).

In P3, probably due to the prolonged lack of interspecies encounters, *A. agrarius* moved towards nocturnality again, becoming cathemeral. Coherently, in P3 its decrease in the percentage of nocturnal visits vs P1 was no longer significant (McNemar’s test: p = 0.219). The mean number of visits of *A. agrarius* in the hours of *A. flavicollis* activity vs the hours of *A. flavicollis* inactivity was again comparable (t = 1.305, p = 0.205), like in P1.

In P4, both species were exposed to an important environmental change. For most mammals, including rodents, the increase in daytime length, and the associated warming up after the cold season, are important triggers of the beginning of the breeding season (Bronson, 2009; Hazlerigg and Simonneaux, 2015; Dardente et al., 2016). In P4 (late winter-early spring), compared to P3, there was an increase in daytime length by over 5500 minutes (P3: 22102 min; P4: 27657 min), with a significant increase of the mean daily daytime length (P3: 650.1; P4: 813.4; t = 14.093, p < 0.0001), which was accompanied by a significant raise of the mean daily maximal temperature by +5.3 °C (P3: 3.1 °C; P4: 8.4 °C; t = 3.629, p = 0.0006), with an increase of +14 °C increase in the lowest daily maximal temperature of the period (P3: −13 °C; P4: +1 °C) and a more than doubled percentage of days with maximal temperature ≥10 °C (P3: 17.6%; P4: 38.2%). In P4, both species increased their percentage of diurnal activity. In particular, the nocturnal activity of *A. agrarius* reached its lowest peak across all 4 time periods: 29.4%. However, while remaining nocturnal, in P4 *A. flavicollis* also increased its percentage of diurnal activity (nocturnal activity dropped to 68.8%), invading the temporal niche of *A. agrarius*. In association with this, *A. agrarius* showed again a significant avoidance of the hours occupied by *A. flavicollis*. In P4, the mean number of visits of *A. agrarius* in the hours when *A. flavicollis* was active was again significantly lower than in the hours when *A. flavicollis* was inactive (t = −2.274, p = 0.033), as observed in P2.

## 4. Discussion

In the present work, we studied the temporal preferences for foraging of two free-living mouse species, *A. agrarius* and *A. flavicollis*, co-inhabiting the same environment. In particular, we examined how the deployment of a novel and highly palatable food source, directly in the natural habitat of these two species, could generate food competition and influence their temporal preferences. In the first period, both species were preferentially nocturnal. However, *A. flavicollis* demonstrated no social tolerance towards *A. agrarius*. When interspecies encounters occurred in the chamber containing the novel food source, *A. flavicollis* showed a very high level of aggressiveness towards *A. agrarius*. In P1, 83.3% of interspecies encounters were agonistic and, within these agonistic interspecies encounters, in 100% of the cases *A. flavicollis* was the attacker and *A. agrarius* was the target of the aggression. Moreover, *A. flavicollis* demonstrated a strong dominance over *A. agrarius*, defeating it in 80% of the cases. Following the interspecies fights of P1, in P2 *A. agrarius*, but not *A. flavicollis*, changed temporal preference, flexibly switching to diurnality and avoiding the hours occupied by *A. flavicollis*. In P3, likely due to the prolonged lack of interspecies fights, *A. agrarius* suspended its hour-specific avoidance behavior and became cathemeral. Nevertheless, in P4, when the environmental changes associated with the onset of the breeding season (increase of daily daytime length and temperature) brought to a relative increase of diurnal activity in both species, *A. agrarius* reacted by resuming a significant avoidance of the hours occupied by *A. flavicollis*, and moved to a strongly diurnal chronotype, while *A. flavicollis* remained nocturnal (as in all previous time periods).

Importantly, in our study we performed a comparative analysis of the behavior of these two sympatric mouse species. According to a comparative approach to the study of behavior (d’Isa and Abramson, 2023; Abramson, 2023), the comparison of the behaviors of two closely related species, especially when they also share the same habitat, provides a unique opportunity to identify species-specific behaviors and, by subtraction of the common potentially causal factors and the common needs, to understand the causes and functions of these species-specific behaviors. In our study, we observed that *A. agrarius* and *A. flavicollis* adopted two very different strategies to obtain access to the novel food source. Indeed, *A. flavicollis* employed physical strength to overpower its competitor, while *A. agrarius* relied on cognitive and behavioral flexibility. More specifically, *A. agrarius* showed clear temporal niche switching when a novel food source was made available in an area that was also visited by the more aggressive and physically dominant *A. flavicollis.* By adjusting its foraging activity to different times of the day or night, *A. agrarius* was able to exploit the new food source when *A. flavicollis* may have been confined to burrows and asleep (Thompson 1982; Johnson and Karels, 2016). This flexibility led to a temporal niche segregation and enabled the species to access food resources that were not exploited by *A. flavicollis*.

Importantly, the temporal niche switching by *A. agrarius* was an effective strategy, as the interspecies encounters showed a dramatic drop in all time periods after P1, while *A. agrarius*’ proportion of visits with successful access to food was increased in all time periods after P1. This flexible temporal niche switching showed by *A. agrarius* could have been achieved through a process of operant conditioning (d’Isa et al., 2011), in which it learned to avoid performing visits within a specific time range after experiencing interspecific attacks and consequent defeat in that time range. Basically, it could be the result of a learned association between a fight and the time of the day when it occurred.

Remarkably, the behavioral flexibility that we observed by *A. agrarius* is coherent with what has been previously reported in the literature for this animal. Indeed, its ability to flexibly adapt to the urban environment is so big that this species has been defined a “urban adapter” (Łopucki and Kiersztyn, 2020; Łopucki et al., 2021). Moreover, it has been found that urban *A. agrarius* mice are better problem solvers than rural *A. agrarius* mice, which suggests that this species may flexibly learn new problem-solving strategies to successfully cope with the complexity of human-altered environments (Dammhahn et al., 2020; Mazza and Guenther, 2021). More recently, our team discovered in the free-living *A. agrarius* a highly flexible dodging behavior (performed to evade a pursuing mouse) which suggests the employment of an intentional tactical deception strategy (d’Isa et al., 2024a).

The nocturnality and the aggressiveness of *A. flavicollis* are also coherent with the previous literature. Indeed, a study of the diel activity of free-living *A. flavicollis* mice in the Białowieża National Park (in north-eastern Poland) found strong and consistent nocturnality by this rodent, as shown by a peak of daily activity in the hours 22.00-2.00, without changes across seasons (Wójcik and Wołk, 1985). In our study, *A. flavicollis* remained consistently nocturnal throughout all four time periods, with a percentage of nocturnality exceeding 87% in the first three periods P1-P3 and still over 68% even after the beginning of the breeding season (in P4).

The natural aggressiveness and dominance of *A. flavicollis* towards the other *Apodemus* species (*A. agrarius* and *A. sylvaticus*) are also well-known (Hoffmeyer, 1973; Montgomery, 1978; Gliwicz, 1981; Mažeikytė, 2002). Moreover, experiments with staged encounters between *A. flavicollis* and *A. sylvaticus* (a mouse of dimension similar to *A. agrarius*) have been performed in the laboratory, finding that *A. flavicollis* was dominant in 90% of the cases (Montgomery, 1978). Our study supports the previous reports that *A. flavicollis* is dominant over *A. agrarius*, as we found that *A. flavicollis* was the winner of the agonistic interactions with *A. agrarius* in 84.6% of the cases.

From a comparative point of view, an interesting question is: after the food competition-related agonistic interactions between the two species, why did *A. agrarius* change its foraging behavior, while *A. flavicollis* did not? An obvious response is that *A. agrarius*, being physically inferior, needed to do so, while *A. flavicollis* did not. *A. flavicollis* is on average larger than *A. agrarius*, at 9-12 cm in body length versus 7-9 cm for the latter. Moreover, behavioral differences also contribute to determine dominance. We have found *A. flavicollis* to be significantly more aggressive than *A. agrarius*, being the initiator of agonism in 100% of the cases. Due to aggressiveness and physical superiority, it is easy for *A. flavicollis* to acquire and defend food resources in the case of interspecific antagonism, as shown by the very high percentage of wins over *A. agrarius* that we recorded.

However, there may also be a second reason for the observed difference in the temporal patterns of the two species. Our chance level analysis revealed that *A. flavicollis* remained consistently nocturnal throughout every period of the study. Interspecies comparisons further showed that *A. flavicollis* was more nocturnal than *A. agrarius* in every period. While *A. agrarius* demonstrated that it can be nocturnal, diurnal or cathemeral (depending on the circumstances), *A. flavicollis* showed only a single chronotype. This consistent nocturnal activity of *A. flavicollis* suggests that the species may be more specialized for operating in low-light conditions. Indeed, the greater relative dimension of the eyes (eyeball to head size ratio) of *A. flavicollis* compared to *A. agrarius* would suggest an anatomical adaptation to exploration and foraging in the dark, as an optimization of the efficiency of light capture, since moonlight is much less intense than sunlight.

Notably, the context in which the outcomes were found (presentation of highly palatable novel food in a peri-urban environment) provides an ideal backdrop to consider whether the species outcompeted in interspecies encounters (*A. agrarius*) may actually be more resilient to environmental changes and more able to survive in hyper-perturbed natural or anthropogenic environments. The rigidity of the activity pattern exhibited by *A. flavicollis* could be attributed to several factors, including physiology of the species, higher efficiency of nocturnal foraging, or being able to avoid predators better when active during the night. Regardless, the behavior of *A. flavicollis* appears far less flexible than *A. agrarius*, potentially exposing it to greater risks in conditions of environmental change. Importantly, researchers have compiled substantial evidence indicating that behavioral plasticity, when present among rodents, confers a distinctive capacity to survive climate change (Rowe et al., 2015; Ramírez-Bautista et al., 2020; Taillie et al., 2023; Hua et al., 2025). From this point of view, *A. agrarius* would certainly be endowed of a greater resilience and adaptability than the physically superior *A. flavicollis*.

## 5. Conclusion

We used a natural field assay to observe interspecies interactions between two species of *Apodemus* mice, finding that under natural conditions, when a highly-palatable food is present, *A. flavicollis* is dominant over *A. agrarius* in the direct interspecific interactions. Nevertheless, after experiencing interspecies fights, *A. agrarius* employs temporal niche switching to segregate its temporal niche from the one of *A. flavicollis*, thereby accessing the food safely, without fights. The flexibility in the activity pattern of *A. agrarius* illustrates a significant adaptive advantage, allowing the species to survive by obtaining food while avoiding a dominant competitor. This behavioral plasticity in switching from nocturnal to diurnal activity is crucial from an ecological perspective, as it allows for coexistence and resource sharing among competing sympatric species that inhabit the same habitat.

Interestingly, rodents are considered the main model system for the study of interspecific competition (Eccard and Ylönen, 2003). The study of interspecific competition between two rodents as *A. agrarius* and *A. flavicollis* not only enhances our understanding of their ecological interactions, but also provides broader insights into the social dynamics and the mechanisms driving species coexistence and community structure. Indeed, from an ecological perspective this expertise can be useful for predicting how animal populations will respond to environmental changes, together with habitat fragmentation, weather alternate, and human interference. On the other hand, from the perspective of social behavior research, this knowledge can contribute to shed light on the strategies that can be adopted by groups interacting with each other in symmetrical or asymmetrical relationships.

Future studies could build on our findings to examine the underlying physiological mechanisms that enable behavioral flexibility and test how the introduction of specific variables affects social behavior and activity patterns. Integrating laboratory-based and naturalistic studies can provide a more comprehensive understanding of biological processes and adaptive behaviors. Such integration can lead to a new paradigm where our understanding of behavioral processes, by taking into account ethologically and ecologically relevant factors, can become more valid, holistic and broadly applicable.

## Ethics statement

This observational study was non-invasive, relying on the monitoring of free-ranging animals that could freely choose to enter or avoid experimental chambers equipped with food and video cameras. As such, it did not require approval from the local ethics committee for animal experimentation. In the present study, no animal was captured nor handled by humans. The observed animals were free-living and remained such during and after the observations. The research was conducted on private property with the landowners’ consent, and all procedures adhered to the Polish Animal Protection Act (August 21, 1997) and the European regulations (Directive 2010/63/EU). The study was designed and implemented in accordance with the ARRIVE guidelines (Kilkenny et al., 2010).

## Data availability statement

Data are available in the supplementary material: complete list of experimental days with dates, civil dawn times, civil dusk times, daytime lengths, nighttime lengths, minimal temperatures and maximal temperatures (Supplementary Table 1); visits per light/dark phase (Supplementary Table 2); visits per hour of the day (Supplementary Table 3); social encounters, all and subcategories (Supplementary Table 4).

