## Supplementary figures and images for "Chronoecological interactions: Temporal niche-switching by black-striped mice after agonistic food competition with a dominant sympatric mouse species"

### Supplementary Figure 1

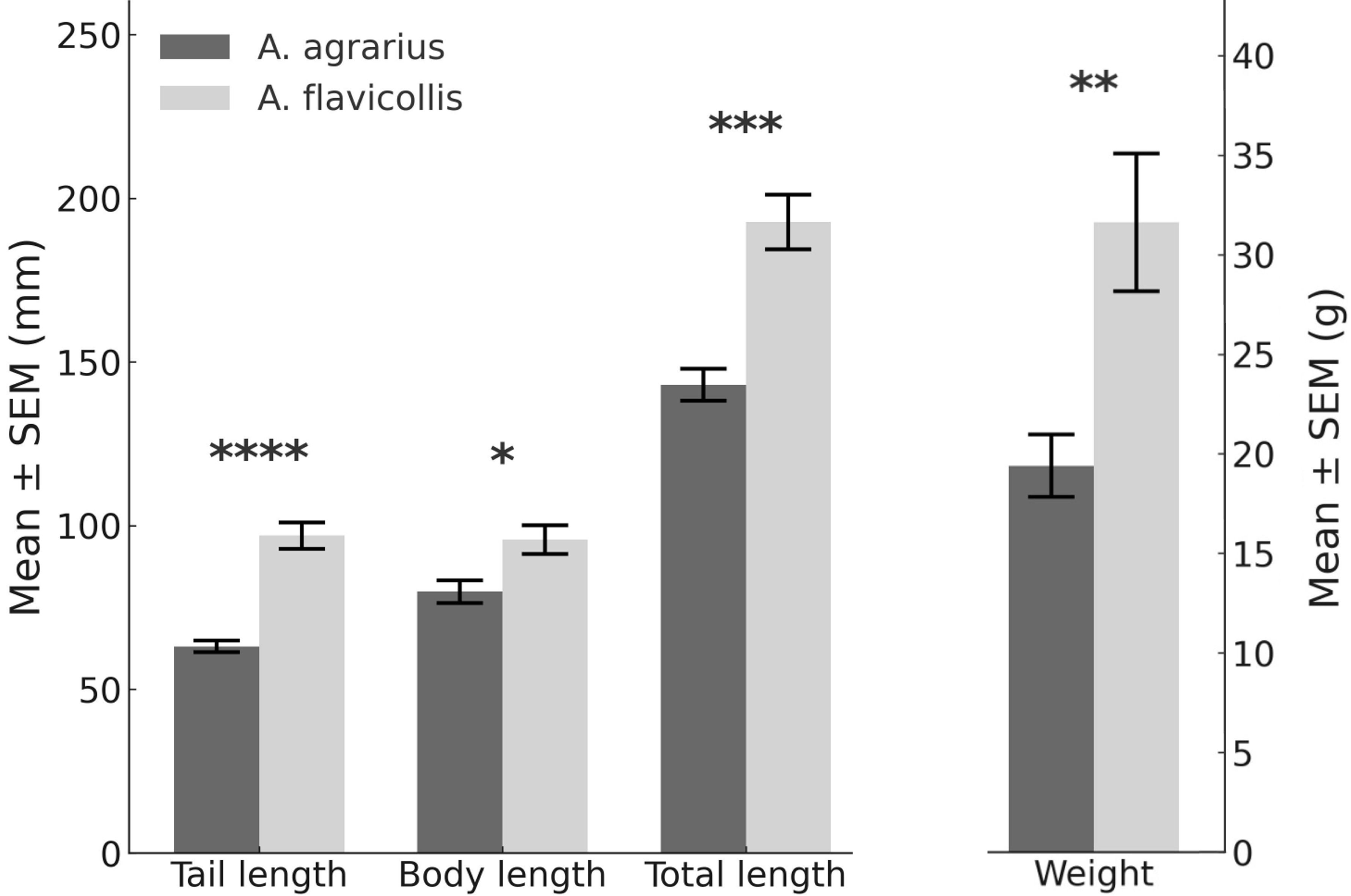
